# Umbilical Cord Blood Cell Transcriptional and Methylation Signatures at Birth Are Associated with BPD Development

**DOI:** 10.1101/2025.11.04.686323

**Authors:** Qianli Liu, Kathryn A. Helmin, Jeanette Bailey, Xóchitl G. Pérez-Leonor, Duc Phan, Hiam Abdala-Valencia, Leena B. Mithal, Benjamin D. Singer, Mary E. Robbins, Marta Perez

**Author notes:** To whom correspondence should be addressed: Marta Perez, MD, Department of Pediatrics/Neonatology, Stanley Manne Children’s Research Institute, 303 E. Superior St., Chicago, IL 60611 USA. Co-senior authors contributed equally to this work (Benjamin D. Singer, Mary E. Robbins, Marta Perez). Competing Interest Statement: BDS holds United States Patent No. US 10,905,706 B2, Compositions and Methods to Accelerate Resolution of Acute Lung Inflammation, and serves on the Scientific Advisory Board of Zoe Biosciences. The other authors have no competing interests to declare.

## Abstract

Bronchopulmonary dysplasia (BPD) is the most common respiratory disease in preterm infants born at less than 28 weeks gestation. Most existing clinical prediction models for BPD show limited accuracy in predicting BPD development when validated using external data, stressing the need for novel biomarkers to identify at-risk infants for early and effective interventions. We leveraged existing frozen umbilical cord blood samples from the Northwestern University Cord Blood Biobank (NUCord) to perform parallel transcriptional and DNA methylation profiling. BPD-associated differentially expressed genes (DEGs) in our cohort included markers previously established in clinical and animal BPD studies, such as genes related to NF-κB signaling and immune responses. We also identified that BPD development is associated with disrupted methylation signatures in microRNA genes and genes associated with glucose metabolism. Our results suggest that BPD development is associated with distinct transcriptomic and epigenetic signatures when compared with healthy term and preterm infants. These signatures may represent biomarkers measurable at birth that predict BPD development during a time window when preventative or therapeutic interventions could be applied.

## Introduction

Bronchopulmonary dysplasia (BPD) remains the most common sequela of premature birth, with an estimated incidence of 10,000–15,000 infants per year in the United States.^1^ Up to 68% of extremely low birth weight (ELBW) infants, defined as infants born at ≤28 weeks of gestation and ≤1000 grams, at National Institute of Child Health and Human Development Neonatal Research Network centers were diagnosed with BPD.^2^ Infants with BPD are at higher risk of mortality and morbidity, including prolonged hospitalization, than age-matched infants without BPD.^3^ In addition, they carry a higher life-long risk of recurrent respiratory exacerbations, cardiovascular complications, growth failure, and neurodevelopmental impairment.^4–6^

The pathophysiology of BPD is multifactorial and influenced by a variety of antenatal and postnatal factors beyond the degree of prematurity.^7,8^ Given the fact that many antenatal factors have been associated with later development of BPD, fetal origins of BPD have been proposed as a potential source of disease pathogenesis, with intrauterine inflammation and oxidative stress identified as likely contributing factors.^9,10^ For example, maternal vascular underperfusion of the placental bed correlates with BPD risk in ELBW infants; moreover, lower expression of cord blood angiogenic factors is associated with higher risk for BPD-associated pulmonary hypertension.^11,12^ Another recent study reported differences in cord blood pathways affecting leukopoiesis and immunity in preterm infants who are later diagnosed with BPD compared to those who are not.^13^

Previous transcriptomic studies of BPD development have predominantly focused on characterizing disease-associated transcriptional and epigenetic signatures in patients with established BPD.^13–17^ However, BPD diagnosis is not made until 36 weeks postmenstrual age, leaving a significant therapeutic window between preterm birth and diagnosis. Most existing clinical prediction models for BPD show limited accuracy in predicting disease development when validated using external data,^18^ highlighting the need for novel biomarkers to identify at-risk infants for early and effective interventions. We observed enhanced expression of DNA methyltransferases, DMNT1, DMNT3A, and DNMT3B, in patients with BPD and in a mouse model that mimics BPD,^19,20^ suggesting that BPD is associated with disrupted methylation signatures.

Whether specific transcriptional and methylation signatures in cord blood could be used as early predictors for BPD development remains unknown. There is a paucity of data regarding developmental changes in transcriptional and methylation signatures present in cord blood between extreme prematurity and term gestation, as well as how these signatures may be altered (or reprogramed) in infants who later develop BPD. Given this knowledge gap, we leveraged the buffy coat of frozen cord blood samples in the Northwestern University Cord Blood Biobank (NUCord) to perform transcriptional and DNA methylation profiling using our published procedures to test the hypothesis that preterm infants who develop BPD have unique DNA methylation and DEG signatures present in their cord blood at the time of birth that are not present in term infants or preterm infants who do not go on to develop BPD.^21–24^

## Results

Demographic and clinical characteristics of the study cohort are summarized in **Table 1**. Infants with Grade 2 or Grade 3^27^ bronchopulmonary dysplasia (BPD, n=3) and those with BPD diagnosed with chorioamnionitis (BPD-c, n=5) were born at lower gestational ages (mean 26 0/7 and 24 0/7 weeks, respectively) and had lower birthweights (mean 738 g and 702 g) compared with preterm controls without chorioamnionitis (Preterm, n=4) and preterm infants with chorioamnionitis (Preterm-c, n=4). Term infants (n=8) and term infants with chorioamnionitis (Term-c, n=6) were delivered at expected gestational ages (38–40 weeks) with average birthweights exceeding 3,400 g. Transcriptional profiling via bulk RNA-sequencing (RNA-seq) was performed on 3–6 samples per group, and methylation profiling via reduced representation bisulfite sequencing (RRBS) was performed for 2–7 samples per group, reflecting specimen availability and quality (**Figure 1**).

**Table 1.**
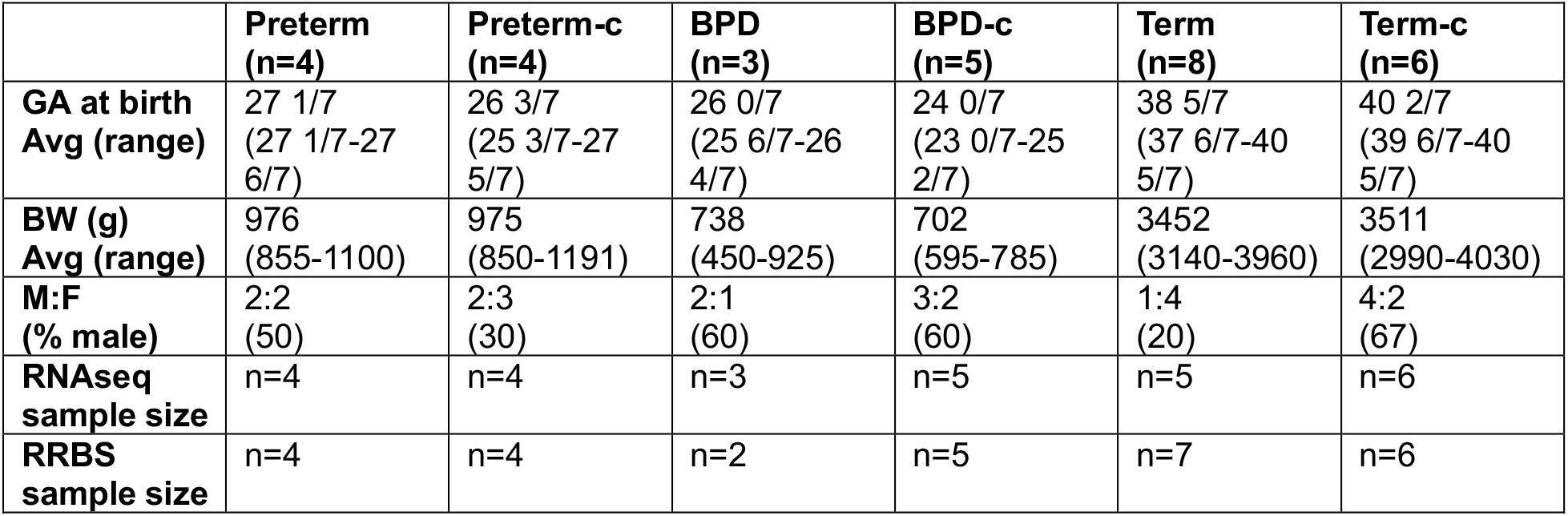
Demographics and clinical information.

**Figure 1.**
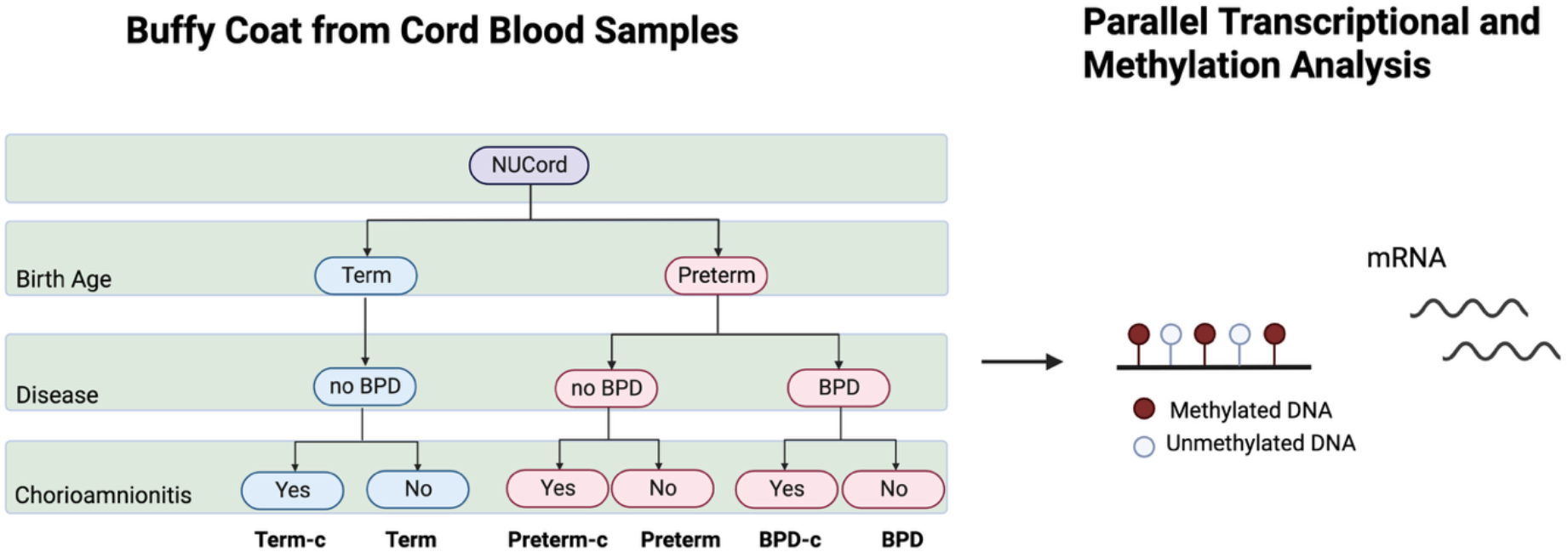
Parallel transcriptional and DNA methylation analysis of umbilical cord blood samples.

We identified 2629 differentially expressed genes (DEGs, false discovery rate [FDR] *q* < 0.05) associated with gestational age and 52 DEGs associated with BPD development (**Figure 2A-B**). For the gestational age comparisons, *k*-means clustering analysis yielded two gene clusters that were then subjected to gene ontology (GO) process and KEGG pathway enrichment analysis. Cluster 1 genes were positively correlated with preterm groups and were enriched in pathways related to oxidative phosphorylation, ATP synthesis coupled electron transport, and the electron transport chain. In contrast, Cluster 2 genes were positively correlated with term groups and were characterized by pathways related to MAPK signaling, chemokine signaling, TNF signaling, and NF-κB-related regulation (**Figure 2A**). Notable DEGs associated with BPD development in our cohort that have been previously identified include: *IL1B* (implicated as an BPD marker in mouse studies^28,29^), *BCL2A1* (downstream target of NF-κB^30^), *ILR2* (involved in mediating inflammatory pathways associated with preterm delivery^31^), *S100A8* (involved in cell cycle progression and differentiation), *CASP10* (involved in regulating inflammatory cell death^32^), *WDFY3* (involved in regulating autophagy^33^), *CD63* (upregulated in the peripheral blood samples from infants with BPD^16^), *NSUN7* (involved in regulating RNA methylation and implicated as a biomarker for neonatal sepsis^34^), and *S100A8* (involved in modulating immune responses^35^) (**Figure 2B-D**). Functional enrichment analysis was not performed for the BPD-specific comparison due to the limited number of DEGs. We also compared the normalized expression of this list of DEGs between all patient groups. In the BPD-associated comparison, we noted that the transcriptional differences among these genes were mainly between the chorioamnionitis and no-chorioamnionitis groups in the preterm groups (**Figure 2B and D**). Nevertheless, this difference was not observed in the age-associated comparison, suggesting that the transcriptional differences were mostly associated with chorioamnionitis infection in preterm infants.

**Figure 2.**
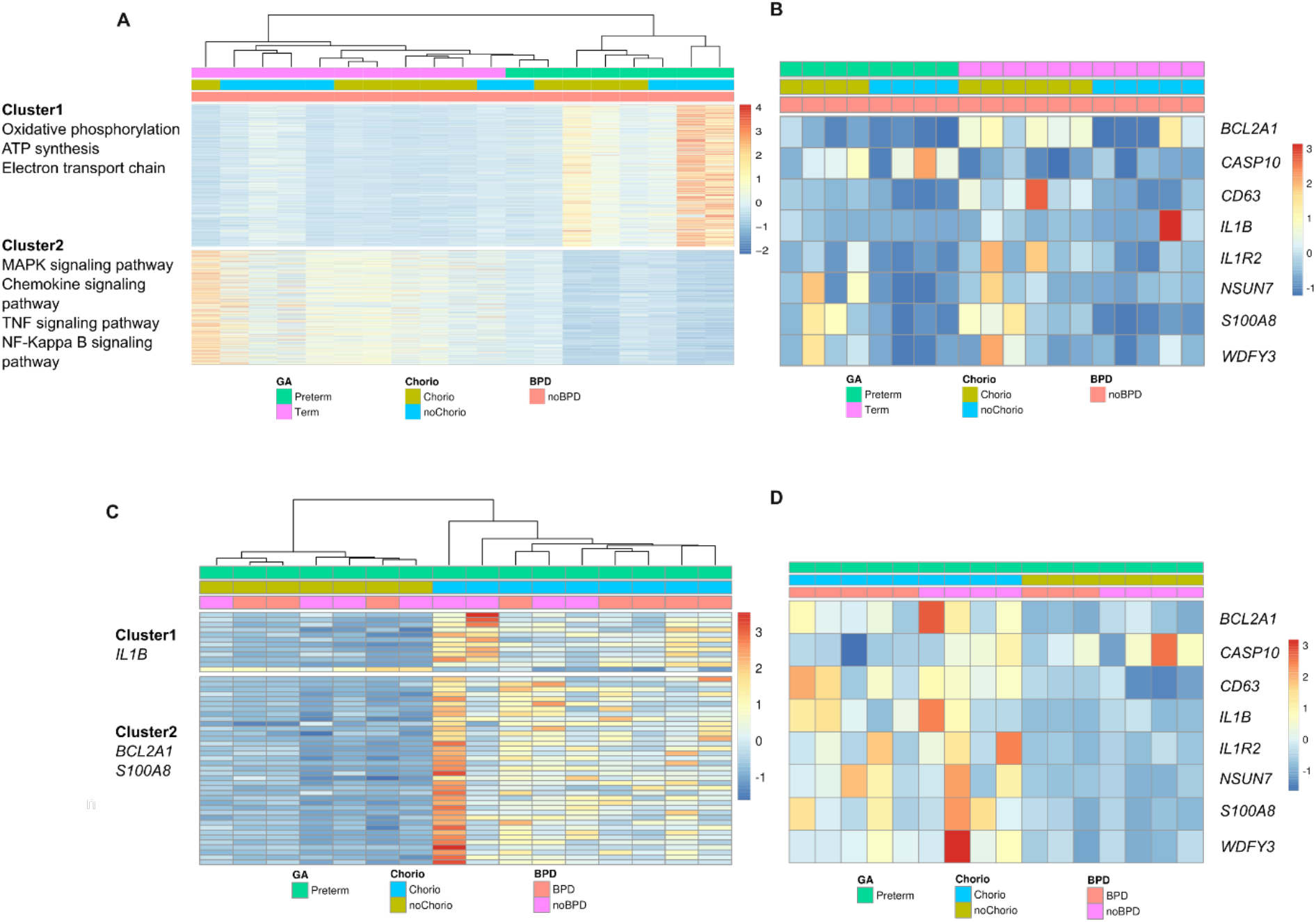
Transcriptional profiling of umbilical cord blood samples. (**A-B**) *k-*means clustering of differentially expressed genes (*q* < 0.05, likelihood-ratio test with FDR correction) from the gestational age analysis (A; n=2629) and BPD analysis (B; n=52). (**C-D**) Expression heatmap of selected genes associated with BPD development from the gestational age analysis (C) and BPD analysis (D).

To investigate whether BPD development is associated with differences in umbilical cord blood immune cell type composition, we leveraged a previously published single-cell RNA-sequencing dataset of peripheral blood mononuclear cells^36^ to perform *in silico* cell-type deconvolution of our bulk RNA-sequencing of umbilical cord blood samples. Major immune cell populations in umbilical cord blood, including NK cells, monocytes, CD4 T cells, CD8 T cells, and B cells, were successfully deconvolved. Our deconvolution results demonstrated that monocytes are the predominant cell type observed across all patient groups (**Figure 3A-B**) and that preterm infants exhibited a lower proportion of monocytes in their umbilical cord blood compared with term infants (**Figure 3A and C**). BPD status did not affect the immune cell composition of the samples (**Figure 3B and D**).

**Figure 3.**
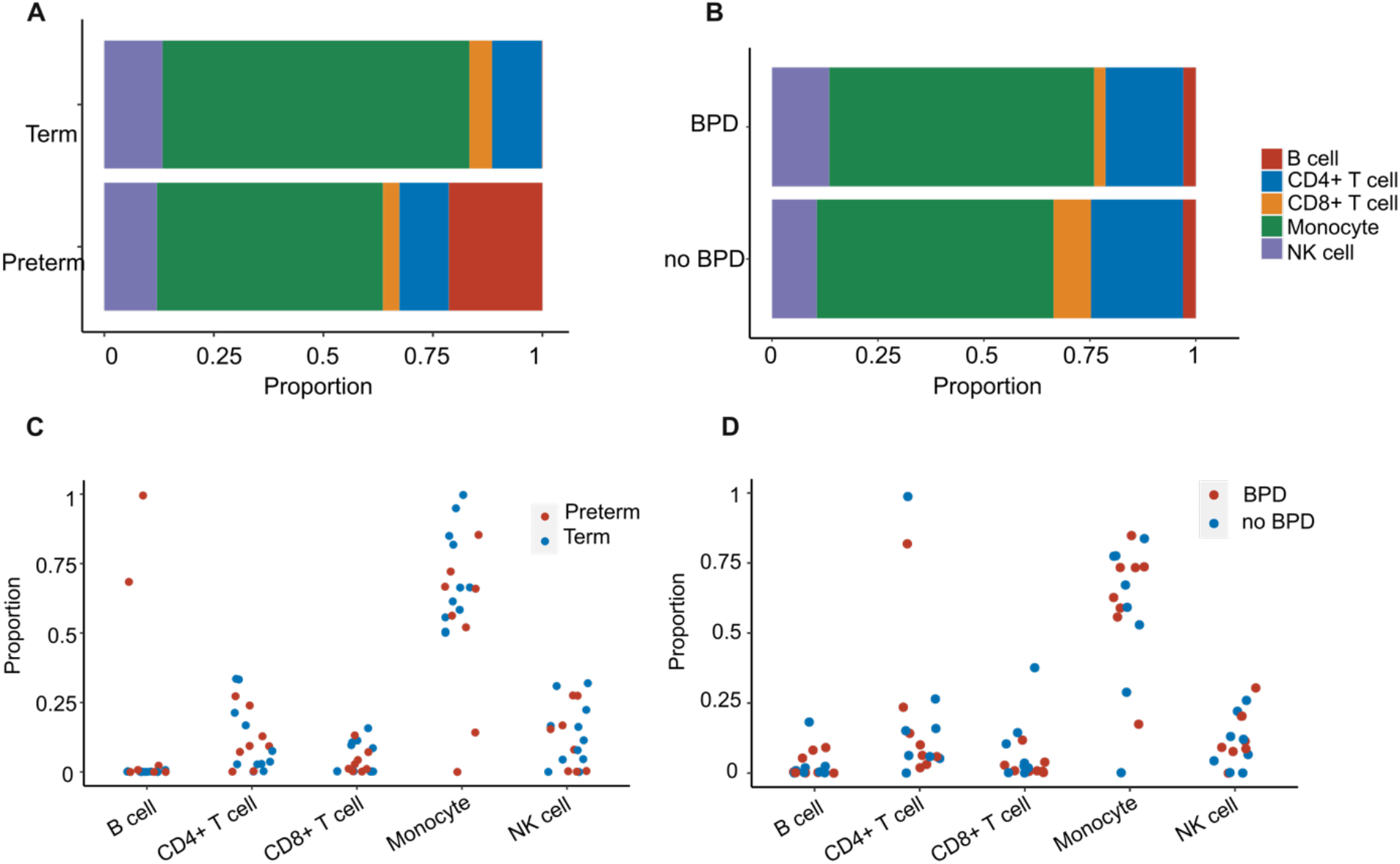
Transcription-based estimation of umbilical cord blood cell type proportion. (**A-D**) Cell type deconvolution analysis showing estimated proportion of B cell, CD4+ T cell, CD8+ T cell, monocyte, and NK cell in the gestational age comparison (A and C) and BPD comparison (B and D).

In parallel with transcriptional profiling, we also performed DNA methylation profiling using our modified reduced-representation bisulfite sequencing (mRRBS) procedure applied to the same cells.^21–23^ We performed differential methylation analysis using Bayesian hierarchical modeling with the Dispersion Shrinkage for Sequencing (DSS) procedure.^37^ Principal component analysis of differentially methylated cytosines (DMCs) with an FDR *q*-value < 0.05 (39,526 for the age-associated comparison and 102,281 DMCs for the BPD-associated comparison) revealed distinct clusters based on BPD development and gestational age (**Figure 4A-B**). The main variance across the BPD-associated comparison corresponded to methylation differences due to BPD development. In the BPD-associated comparison, umbilical cord blood samples from the BPD group were hypermethylated compared with all other groups (**Figure 4C**). In the age-associated comparison, differences in methylation status were mostly associated with gestational age, consistent with the transcriptional results (**Figure 4D**). We also identified genes containing DMCs in their promoter regions in the age-associated and BPD-associated comparisons (**Figure 4E-F**). *K*-means clustering of these genes in the age-associated comparison demonstrated that Cluster 1 genes are hypomethylated in preterm infants; these genes are associated with neonatal development (*FBXO40, TM4SF1*). For the BPD-associated comparison, Cluster 4 genes were hypomethylated in BPD group and include microRNA genes (*MIR6854, MIR3941, MIR6748, MIR140*) and genes associated with disrupted glucose metabolism (*SLC2A3p2*).

**Figure 4.**
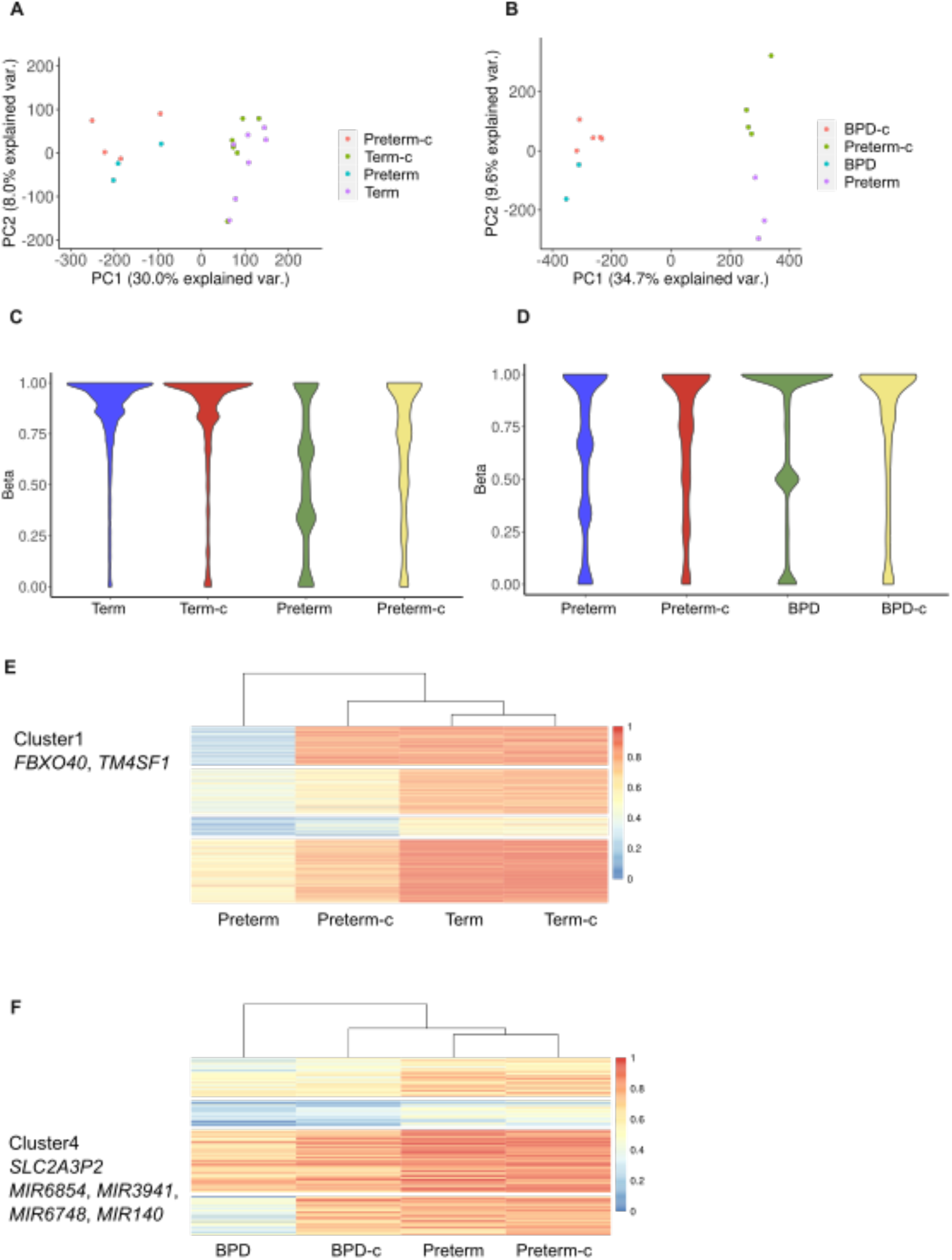
GA and BPD development are associated with distinct DNA methylation signatures. (**A-B**) Principal component analysis of differentially methylated cytosines (DMCs) identified from a linear model and ANOVA-like testing with FDR *q* < 0.05 in the gestational age comparison (A, n=39,526) and the BPD comparison (B, n=102,281). (**C-D**) Violin plots of DMCs distribution in the gestational age comparison (C), and the BPD comparison (D). (**E-F**) *K*-means clustering of gene promoters including DMCs with a difference of ≥20% in methylation between preterm and term infants (E) and between infants who later developed BPD and preterm controls (F). Promoters were defined as 1 kb flanking the transcriptional start sites (TSSs).

## Discussion

Despite substantial advances in neonatal care, the incidence of BPD has largely remained unchanged, reflecting increased survival of extremely preterm infants.^38^ Due to the fact that the diagnosis of BPD is not made until 36 weeks of postmenstrual age, it remains challenging to identify at-risk infants at the time of birth who would benefit from early intervention and treatment. In our parallel transcriptional and DNA methylation profiling of umbilical cord blood, we found that infants exposed to chorioamnionitis who later developed BPD exhibited greater expression of *IL1B* and *BCL2A1*, which are associated with immune activation and pro-inflammatory responses.^28–30^ These results are consistent with previous findings that increased inflammation in extremely preterm infants at birth is correlated with later BPD development.^39^ Our results also demonstrate that BPD development is associated with distinct DNA methylation patterns at birth. We found that while BPD development explains the most variance in methylation level among the four preterm groups (Preterm, Preterm-c, BPD and BPD-c), these preterm samples clustered better based on the chorioamnionitis infection status using the differentially expressed genes. These observations suggest that DNA methylation may be an age-related modifier of the host response to infection and subsequent risk of BPD.

BPD pathogenesis is often described as a developmental growth arrest in the lungs that alters the transition from the canalicular to saccular phases in ELBW infants;^40^ therefore, it was important to consider the fetal developmental trajectory as a variable in our study. From a developmental perspective, our transcriptional data showing differences in immune cell activation between term and preterm infants are not surprising, as premature infants are known to have a blunted immune response compared with full term peers.^41^ Our results also show an greater expression of genes associated with NF-kB pathways in cord blood samples from term infants with chorioamnionitis, which is consistent with a previous study investigating cord blood monocytes’ response to bacterial infection.^42^ These alterations in the NF-kB signal cascade, which has been associated with BPD development in human and mouse models,^43^ could reflect an altered developmental transition. Consistent with previous studies,^44,45^ our methylation data also highlight that there are distinct methylation patterns associated with gestational age. Our results demonstrate that cord blood samples from preterm infants are hypomethylated compared with cord blood samples from term infants when assessing all differentially methylated cytosines and genes associated with neonatal development.

Based on our deconvolution analysis, we were able to determine that monocytes were the predominant cell population in our samples; therefore, the transcriptional and methylation signatures identified should be interpreted from a myeloid cell-response perspective. Not surprisingly, in our cohort, the transcriptional signal associated with chorioamnionitis was more robust than the one associated with BPD diagnosis. In future cord blood studies, investigations may need to pursue isolation of other RNA-containing components in circulation, such as macrovesicles or microvesicles, to identify immune-independent signatures predictive of BPD development.

We previously identified epigenetic silencing of microRNA cluster 17-92 (miR-17∼92) expression in autopsy lung samples from patients with BPD as well as lower serum concentrations of cluster components in the first week of life for survivors who later developed BPD.^19^ Nevertheless, we found no significant differences in miR-17∼92 expression and methylation status in umbilical cord blood samples between healthy infants and infants who later developed BPD. These results suggest that miR-17∼92 is a lung-specific BPD biomarker or a delayed biomarker for BPD development that is not present in umbilical cord blood at birth. Our *k*-means clustering of gene promoters, including DMCs with a difference of ≥20% in methylation between BPD and non-BPD groups, identified additional microRNAs that are hypomethylated in their promoter regions and were found in umbilical cord blood. Prospective studies are needed to validate these features as predictive biomarkers.

Due to the limited availability of cord blood samples from infants who later developed grade 2 or grade 3 BPD, our study is limited by the small sample size in the BPD groups. We also noted substantial transcriptional heterogeneity within the patient groups who later developed BPD (BPD and BPD-c), which further limits the power of our statistical analysis. Overall, our study demonstrates that BPD development combined with chorioamnionitis infection is associated with distinct transcriptional and DNA methylation signatures. Examining a larger cohort of infants who later develop BPD, and possibly a different component of the cord blood, is necessary to further investigate BPD-specific molecular markers in cord blood that could be used to identify infants at highest risk of BPD development.

## Methods

### Cord blood collection and processing

All patients were enrolled in Prentice Birth Cohort within the NUCord biorepository under Northwestern University IRB STU00201858 and Lurie Children’s Hospital IRB 2018–2145. Umbilical cord blood samples were collected at the time of birth, processed using our published methods^46^, and stored at -80 °C. DNA and RNA were extracted from the buffy coat cell pellets using the AllPrep DNA/RNA Micro Kit (QIAGEN).

### RNA-sequencing and analysis

RNA-seq libraries were prepared using the SMARTer Stranded Total RNA-Seq Kit, version 2 (Takara cat. no. 634411) and sequenced using a NextSeq 2000 instrument (Illumina). Analysis was performed using previously published procedures.^47^ In short, after demultiplexing using bcl-Convert (version 3.10.5; Illumina), adaptor trimming, alignment to the GRCh38 reference genome, and quantification were performed using the nf-core/RNA-seq pipeline version 3.9 (implemented in Nextflow 22.04.5 with Northwestern University Genomics Compute Cluster configuration (nextflow run nf-core/rnaseq -profile nu_genomics --genome GRCm38). Samples with less than 5% uniquely mapped reads were excluded from downstream analysis. Differential expression analysis was performed in R package DEseq2 (version 1.38.3 in R 4.2.3). Differentially expressed genes (FDR *q* < 0.05) were used for *k*-means clustering analysis and *k* was determined by the elbow method. GO term enrichment analysis was performed using R package topGO version 2.50.0 with Fisher’s exact test.

### Modified reduced representation bisulfite sequencing (mRRBS) and analysis

mRRBS was prepared using our custom workflow.^21,23,47^ DNA methylation library quality was assessed using high-sensitivity screen tape (TapeStation 4200, Agilent). 4–6 libraries were pooled with equal molarity and then sequenced using a NextSeq 2000 instrument (Illumina; 75 base-pair singe-end reads). DNA methylation analysis was performed using our previously published Bismark-based pipelines.^22^

### Cell type deconvolution of RNA-sequencing

Cell type deconvolution of the bulk RNA-sequencing analysis was performed using python package cellanneal.^48^ A previously published single-cell RNA analysis of peripheral blood mononuclear cells^36^ was used to derive gene signatures for the deconvolution. Gene dictionary was first generated, and cell type deconvolution was performed using the cellanneal.deconvolve function (maxiter=750).

### Statistical analysis

All statistical analysis were performed using R 4.2.3 and GraphPad Prism version 10.3.1.

## Funding & Acknowledgements

QL is supported by the David W. Cugell Fellowship and the Genomics Network (GeNe) Pilot Project Funding. BDS is supported by NIH awards R01HL149883, R01HL153122, P01HL154998, P01AG049665, U19AI135964, and U19AI181102. We wish to acknowledge the Northwestern University Flow Cytometry Core Facility supported by CA060553; the BD FACSAria SORP system was purchased with the support of S10OD011996. We also wish to acknowledge the Northwestern University Metabolomics and Integrative Genomics Core. This research was supported in part through the computational resources and staff contributions provided by the Genomics Compute Cluster, which is jointly supported by the Feinberg School of Medicine, the Center for Genetic Medicine, and Feinberg’s Department of Biochemistry and Molecular Genetics, the Office of the Provost, the Office for Research, and Northwestern Information Technology. The Genomics Compute Cluster is part of Quest, Northwestern University’s high performance computing facility, with the purpose to advance research in genomics. Figure 1 was created using biorender.com.

## Author Contributions

QL contributed to the conceptualization and methodology of this work; the data generation and visualization; and to the writing and editing of the manuscript. KAH contributed to the methodology, generation of data, and the editing of the manuscript. JB contributed to the generation of data and the editing of the manuscript. XGP-L contributed to the methodology, generation of data, and the editing of the manuscript. DP contributed to the methodology, generation of data, and the editing of the manuscript. HA-V contributed to the methodology, generation of data, and the editing of the manuscript. LBM contributed to the methodology, and the editing of the manuscript. BDS contributed to the conceptualization and methodology of this work; the data generation and visualization; the supervision, administration, and funding acquisition of this work; and to the writing and editing of the manuscript. MER contributed to the conceptualization and methodology of this work; the data generation and visualization; the supervision, administration, and funding acquisition of this work; and to the writing and editing of the manuscript. MP contributed to the conceptualization and methodology of this work; the data generation and visualization; the supervision, administration, and funding acquisition of this work; and to the writing and editing of the manuscript.

